# A database for large-scale docking and experimental results

**DOI:** 10.1101/2025.02.25.639879

**Authors:** Brendan W. Hall, Tia A. Tummino, Khanh Tang, John J. Irwin, Brian K. Shoichet

**Author notes:** These authors contributed equally (Brendan W. Hall, Tia A. Tummino).

## Abstract

The rapid expansion of readily accessible compounds over the past six years has transformed molecular docking, improving hit rates and affinities. While many millions of molecules may score well in a docking campaign, the results are rarely fully shared, hindering the benchmarking of machine learning and chemical space exploration methods that seek to explore the expanding chemical spaces. To address this gap, we develop a website providing access to recent large library campaigns, including poses, scores, and *in vitro* results for campaigns against 11 targets, with 6.3 billion molecules docked and 3729 compounds experimentally tested. In a simple proof-of-concept study that speaks to the new library’s utility, we use the new database to train machine learning models to predict docking scores and to find the top 0.01% scoring molecules while evaluating only 1% of the library. Even in these proof-of-concept studies, some interesting trends emerge: unsurprisingly, as models train on larger sets, they perform better; less expected, models could achieve high correlations with docking scores and yet still fail to enrich the new docking-discovered ligands, or even the top 0.01% of docking-ranked molecules. It will be interesting to see how these trends develop for methods more sophisticated than the simple proof-of-concept studies undertaken here; the database is openly available at lsd.docking.org.

## Introduction

In the last six years, with the advent of make-on-demand (“tangible”) libraries^1,2^, readily available chemical space has increased by over four orders of magnitude. Structural modeling of these libraries has supported a concomitant growth in the size and success of prospective molecular docking campaigns^3–8^, with applications ranging from G protein-coupled receptors (GPCRs)^5, 9, 10^, to soluble enzymes^11, 12^, transporters^7^, and other targets. The first screens of these tangible libraries consisted of only 99 to 138 million molecules^1^ and have grown to over 1 billion explicitly docked molecules^9, 13^ more recently. Even larger docking campaigns exceeding 11 billion interpolated synthons^14^ and adaptively sampling from 69 billion molecules have been undertaken^15^. While most of these campaigns typically test fewer than 100 docking-prioritized molecules from the larger libraries, for several targets hundreds of molecules have been tested, and overall the number of tested predictions from large-scale docking (LSD) exceeds 10,000 molecules in the open literature^16–24^.

The results of these campaigns, both experimental and simply docking scores and poses, could benefit the community, acting as benchmarking and training sets for machine learning (ML) among other applications^25–36^. However, these results have not been made available at scale in a readily accessible way. For instance, a large library docking paper may report docking scores only for the synthesized molecules^11, 37, 38^ or sometimes only for those that were experimentally active^7, 12^, representing a tiny portion of the overall docked library or even its high-ranking subset. While some papers report scores for many of the docked molecules, hosted on open-source platforms like GitHub^8^ or other publicly accessible websites^1, 10, 39^, information beyond the docking scores, such as docked poses or the energy potential grids used to score the molecules, is rarely provided.

Here we describe an open-source benchmarking set of published docking results for 6.3 billion explicitly evaluated molecules across eleven protein targets (lsd.docking.org). The benchmarking set includes docking scores (using DOCK3.7/3.8), SMILES, poses for top molecules, results for *in vitro* tested molecules, and the docking energy grids used to produce them. As a simple proof of concept, we consider the ability of ML models trained on this set with the widely used Chemprop framework^40^ to predict docking scores (subsequently referred to as Chemprop models). Applications of these models to the chemical space exploration technique Retrieval Augmented Docking (RAD)^41^ are considered.

## Results

### Organization of large-library docking data

We began by collecting the source files for the published large-scale docking campaigns run in our lab to date, covering 11 different targets (**Table 1**, **Fig. 1A, Table S1**). Within a given target, one or more separate large-scale screens were run, typically differentiated by ligand size (e.g., lead-like versus fragments), docking grids (e.g., input structure differences or different parameter optimizations), or purchasability (e.g., in-stock versus make-on-demand).

**Figure 1.**
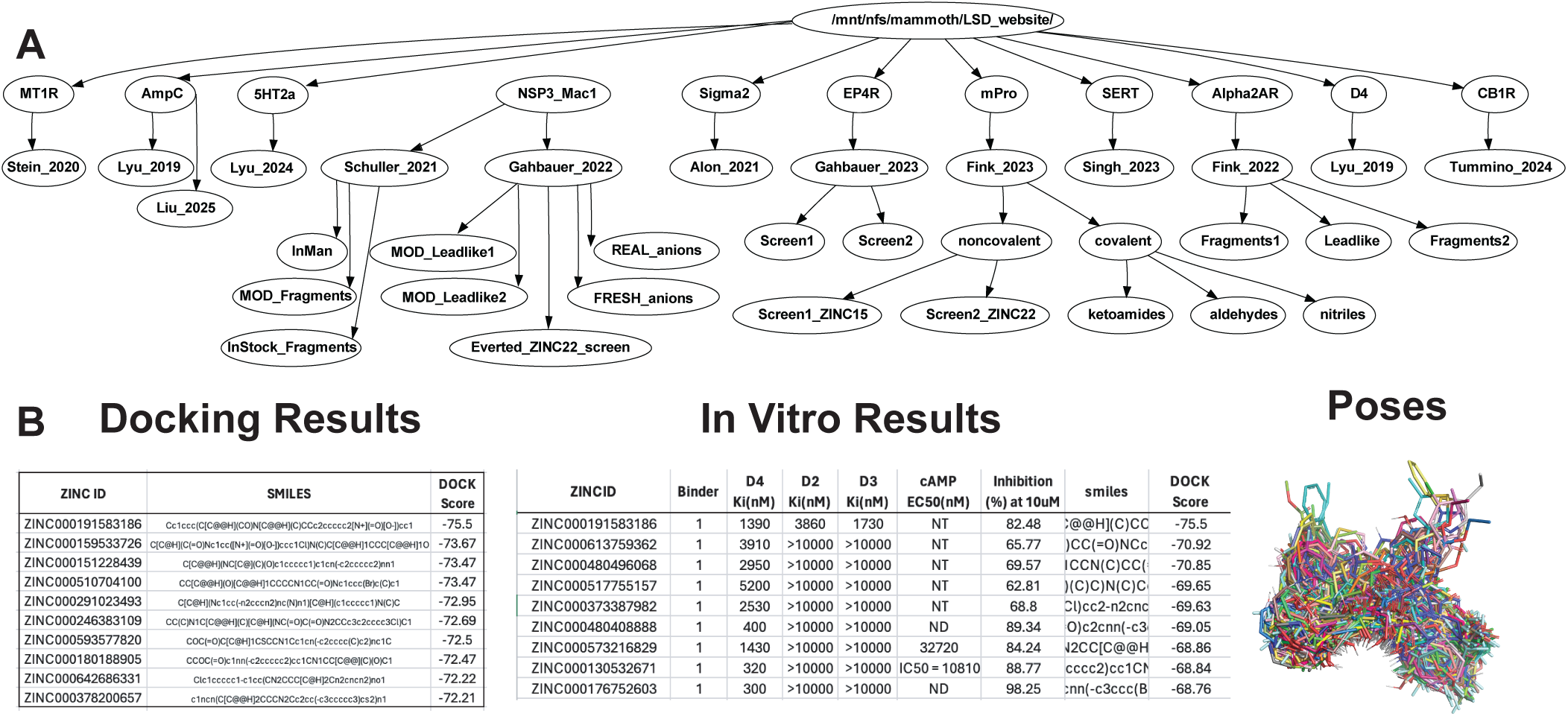
**(A)** Overview of the directory structure on lsd.docking.org. **(B)** Types of data available for each screen: docking scores, *in vitro* experimental data, and poses for the best-scoring molecules.

**Table 1.**
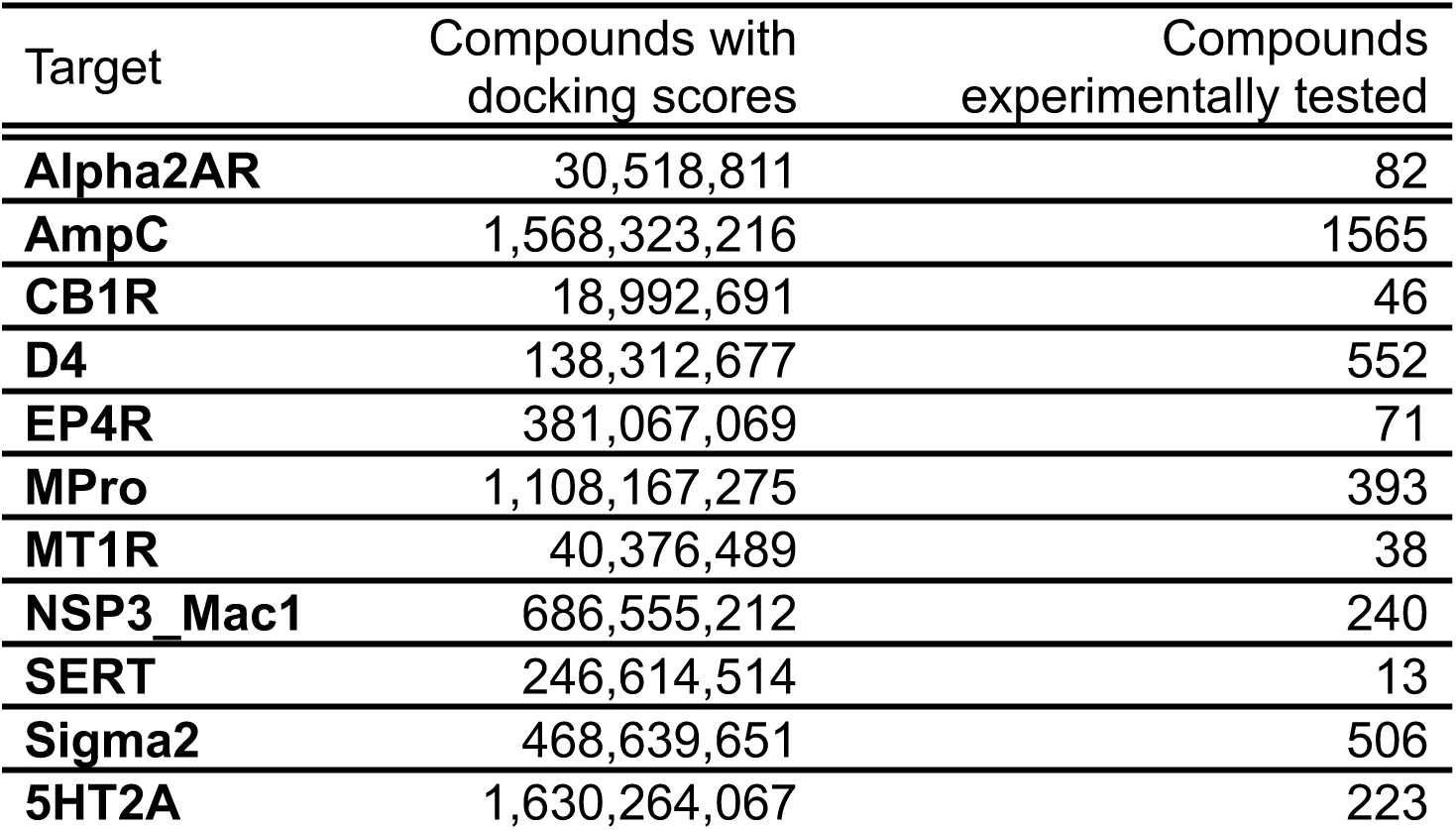
Overview of the docking screens included on lsd.docking.org and the number of docking scores and experimental results provided for each target.

For each screen, we gathered three levels of data (**Fig. 1B**). First, we collected what may be the most useful to the ML community, the “Docking Results”. These include the list of scored molecules, their SMILES, and the docking score for each molecule from every screen. To support these, we also collected the docking energy potential grids used to score the molecules in each screen.

Next, we collected 3D poses for the top 500,000 unfiltered molecules from each screen in mol2 format, which can be visualized with software such as Chimera^42^, PyMoL^43^, or Maestro^44^, among others. Finally, we collected the *in vitro* experimental results and merged them with the docking results for *in vitro* tested molecules. Together, the set provides docking scores for over 6.3 billion ligand-target pairs and provides over 10 million docking poses for the high-ranking molecules from each docking campaign.

To make these data readily accessible, we developed the website lsd.docking.org. Because some targets have been the focus of multiple large-library screens, and because some papers screened multiple targets, users can search either by target or by paper to access the data. Each level of data, docking results, poses, *in vitro* results, and docking grids (dockfiles), is a subfolder nested within that of the individual screen. If multiple files are hosted within a folder, a zip file containing the contents is provided for easier dissemination. A README.md is provided within each screen folder explaining the contents of the subfolders.

For example, to access the docking scores for the dopamine D4 receptor screen, a user would click the “Browse by Target” button which navigates to a page showing every available target. Clicking “View” for the D4 target shows a folder called “Lyu_2019” and clicking the folder brings the user to a page showing the folders for each level of data from the screen: “dockfiles,” “Poses,” “InVitroResults,” and “DockingResults.” Finally, clicking on “DockingResults” navigates to a screen with the downloadable file “D4_screen_table.csv.gz” which contains the ZINC IDs, SMILES, and docking scores for every molecule in the screen.

### Impact of training set size and sampling on Chemprop performance

While we do not introduce or apply new ML methods here, we thought it would be interesting to train ML models with these datasets as a proof of concept for the possible utility of this dataset. Accordingly, we applied the well-known framework Chemprop^40, 45^ on subsets of the docking scores from campaigns against AmpC β-lactamase (AmpC)^4^, 5HT2A^9^, and Sigma2^39^, containing 1,468,863,655, 1,630,264,067, and 468,639,651 molecules, respectively. The large datasets allowed us to systematically investigate the effects of training set size and data sampling strategies.

We investigated training set sizes of 1,000, 10,000, 100,000, and 1,000,000 molecules sampled from the datasets using three strategies: random sampling across the entire dataset, random sampling from the top ranking 1% of molecules only, and a stratified approach where 80% of the training set was randomly sampled from the top ranking 1% of molecules and the remaining 20% was sampled from the rest of the dataset. We created test sets by randomly sampling 100,000,000 molecules from each dataset. We evaluated model performance using the Pearson correlation between predicted and true scores across the entire test set and the top 0.01% scoring molecules, ensuring no overlap with the training set. Additionally, we measured the logAUC, which quantifies the fraction of the top 0.01% molecules found as a function of the screened library fraction on the logarithmic scale (**Fig. 2C**). This metric captures the model’s ability to enrich for the true top 0.01% scoring molecules in its top predictions and is widely used in docking studies^46–48^.

**Figure 2.**
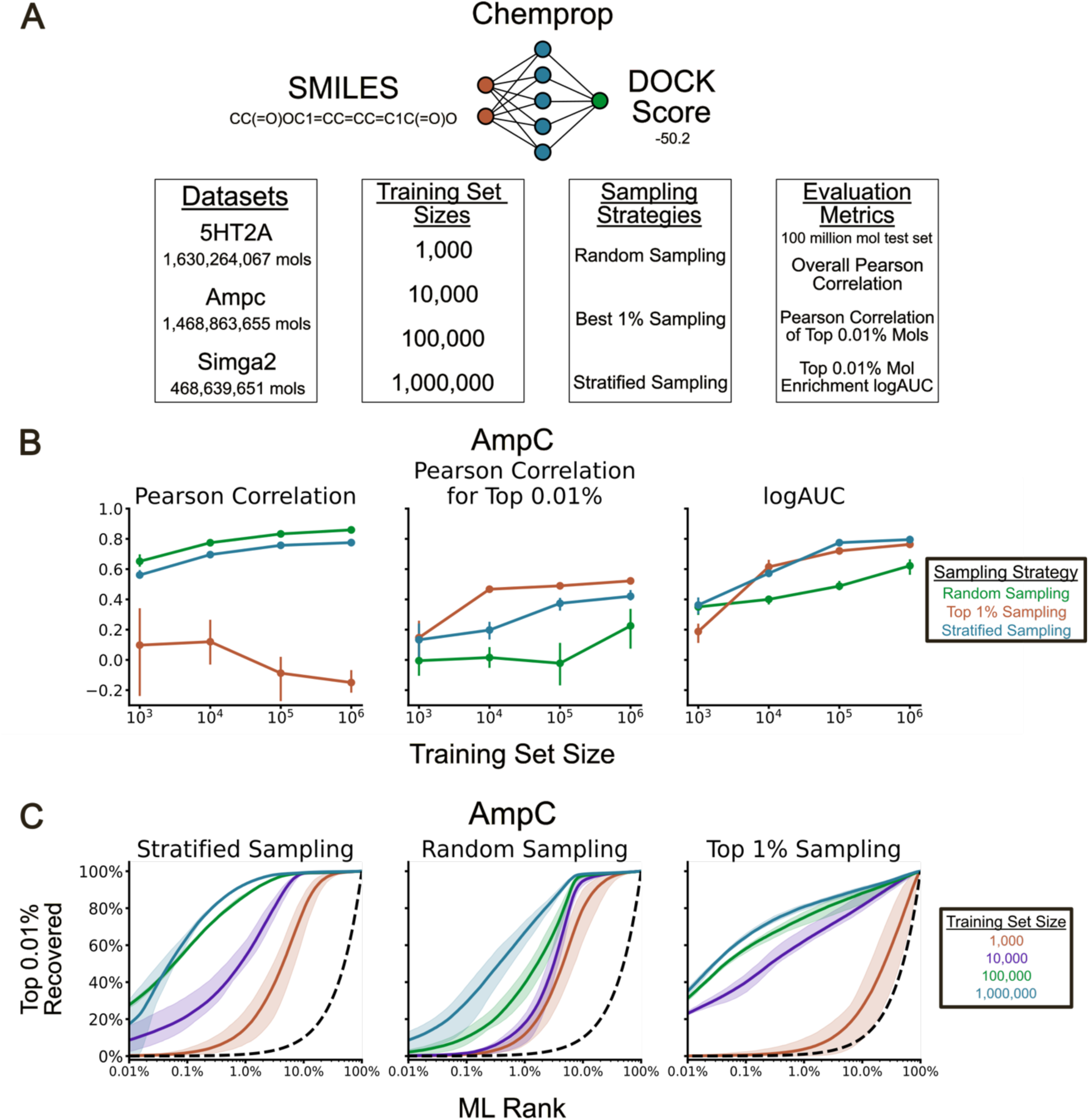
**(A)** Overview of the datasets, training set sizes, sampling strategies, and evaluation metrics used to investigate the ability of Chemprop models to predict DOCK scores. **(B)** Effects of increasing the training set size on the three evaluation metrics for AmpC. The bars represent the minimum and maximum performance across five replicates. **(C)** AmpC log recall curves of the top 0.01% scoring molecules for the three sampling strategies using increasing training set sizes. The shaded region represents the minimum and maximum performance across five replicates. The dashed black line represents a random predictor.

Unsurprisingly, increasing the training set size improved model performance across nearly all targets, sampling methods, and evaluation metrics (**Fig. 2B**, **Fig. 2C**). For instance, AmpC models trained with random sampling achieved overall Pearson correlations of 0.65, 0.77, 0.83, and 0.86 between predicted and true scores when trained with 1,000, 10,000, 100,000, and 1,000,000 molecules, respectively (**Fig. 2B**) (we note that even a million molecules are only 0.07% of the overall library docked). Intriguingly, the overall Pearson correlation between a model’s predictions and true scores did not reliably indicate its ability to enrich for the top 0.01% scoring molecules or for the true binders. For example, on the AmpC dataset with 100,000 training molecules (0.007% of the docked library), random sampling achieved a Pearson correlation of 0.83, higher than the 0.76 with stratified sampling (**Fig. 2B**). However, the same random sampling achieved a logAUC of only 0.49 for the recall of the top 0.01% scoring molecules, whereas stratified sampling achieved a logAUC of 0.77 for this 0.01%--i.e., many more of the highest-ranking molecules were found in the latter’s top predictions than the former’s, despite the former’s better *R-*value. Similarly, random sampling only achieved a logAUC of 0.47 for the recall of true, novel inhibitors found in the AmpC docking campaign, whereas stratified sampling had a logAUC of 0.87 for these inhibitors (**Fig. 2C and Fig. 3B**).

**Figure 3.**
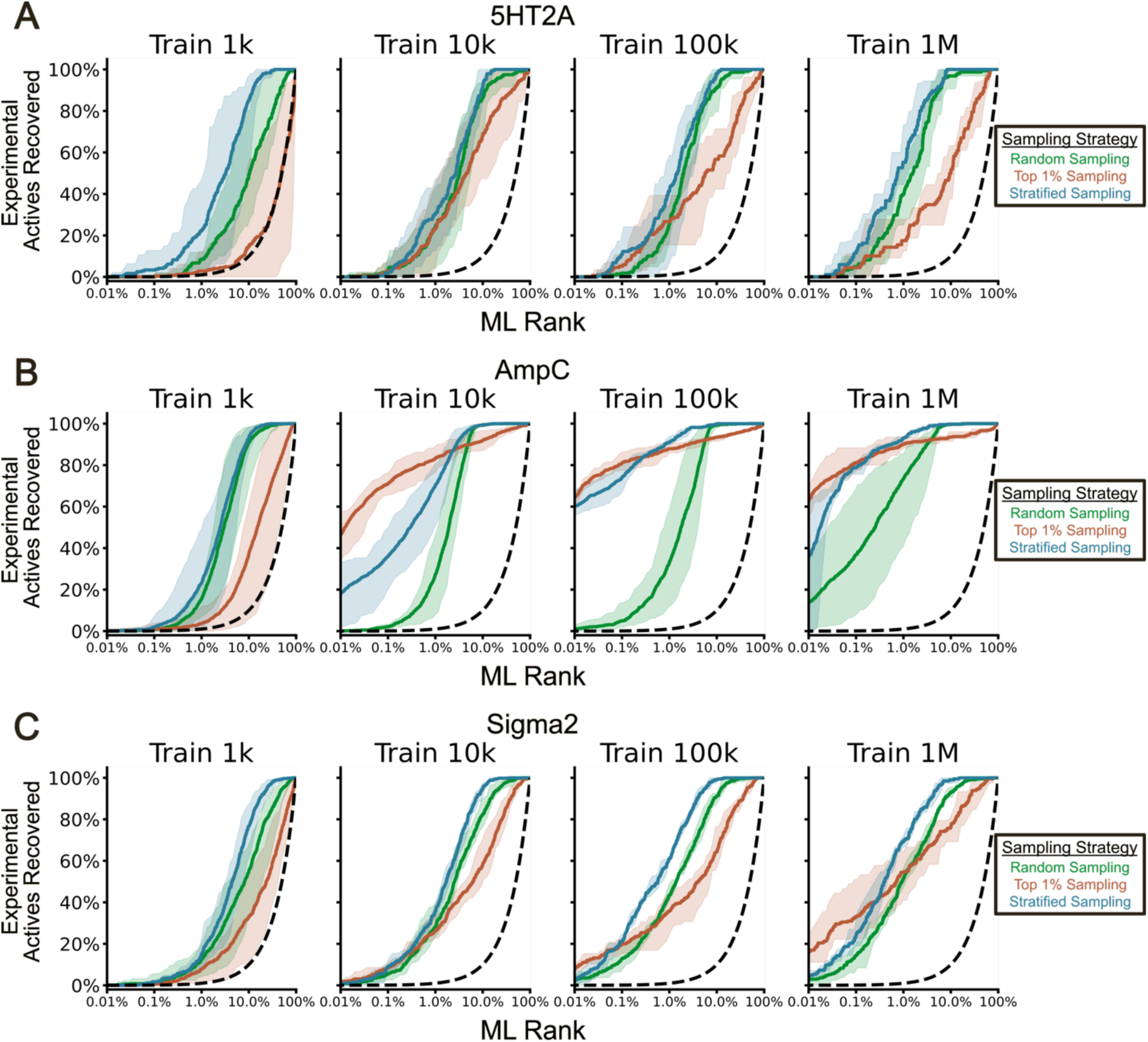
Chemprop model recall curves of the experimental actives after filtering out molecules that share a scaffold with any molecule in the training set for the **(A)** 5HT2A receptor, **(B)** AmpC receptor, and **(C)** Sigma2 receptor. The shaded region represents the minimum and maximum performance across five replicates. The dashed black line represents a random predictor.

Overall, despite relatively high Pearson correlations, few of the top 0.01% scoring molecules and the true ligands were in the top 0.01% of ML predictions – the region realistically inspected during large-scale screening. For example, the random sampling strategy on the AmpC dataset with 1 million training molecules (0.05% of the docked library) achieved the highest Pearson correlation of 0.86, but only 8.4% of the top 0.01% scoring molecules and 13.5% of the true binders were in its top 0.01% predictions. These results also held true for the 5HT2A and Sigma datasets (**Fig. 3, Fig. S1, Fig. S2**).

### Retrieval Augmented Docking using Chemprop models

We next investigated the utility of the lsd.docking.org benchmarks for evaluating chemical space exploration methods. We assessed the ability of Retrieval Augmented Docking (RAD)^41^ to find the top 0.01% scoring molecules from billion-scale chemical libraries. Drawing on a technique from large-scale vector databases, as used in social networks, RAD organizes chemical libraries into Hierarchical Navigable Small World (HNSW) graphs^49^, enabling efficient exploration of chemical space (**Fig. 4A**). Previous studies found that traversing the HNSW graph with the DOCK3.7 scoring function finds most of the top 0.01% scoring molecules while only needing to explicitly evaluate a fraction of the library. Here, we constructed HNSW graphs for ∼1.3 billion molecules from the AmpC and 5HT2A datasets, and ∼500 million molecules from the Sigma2 dataset.

**Figure 4.**
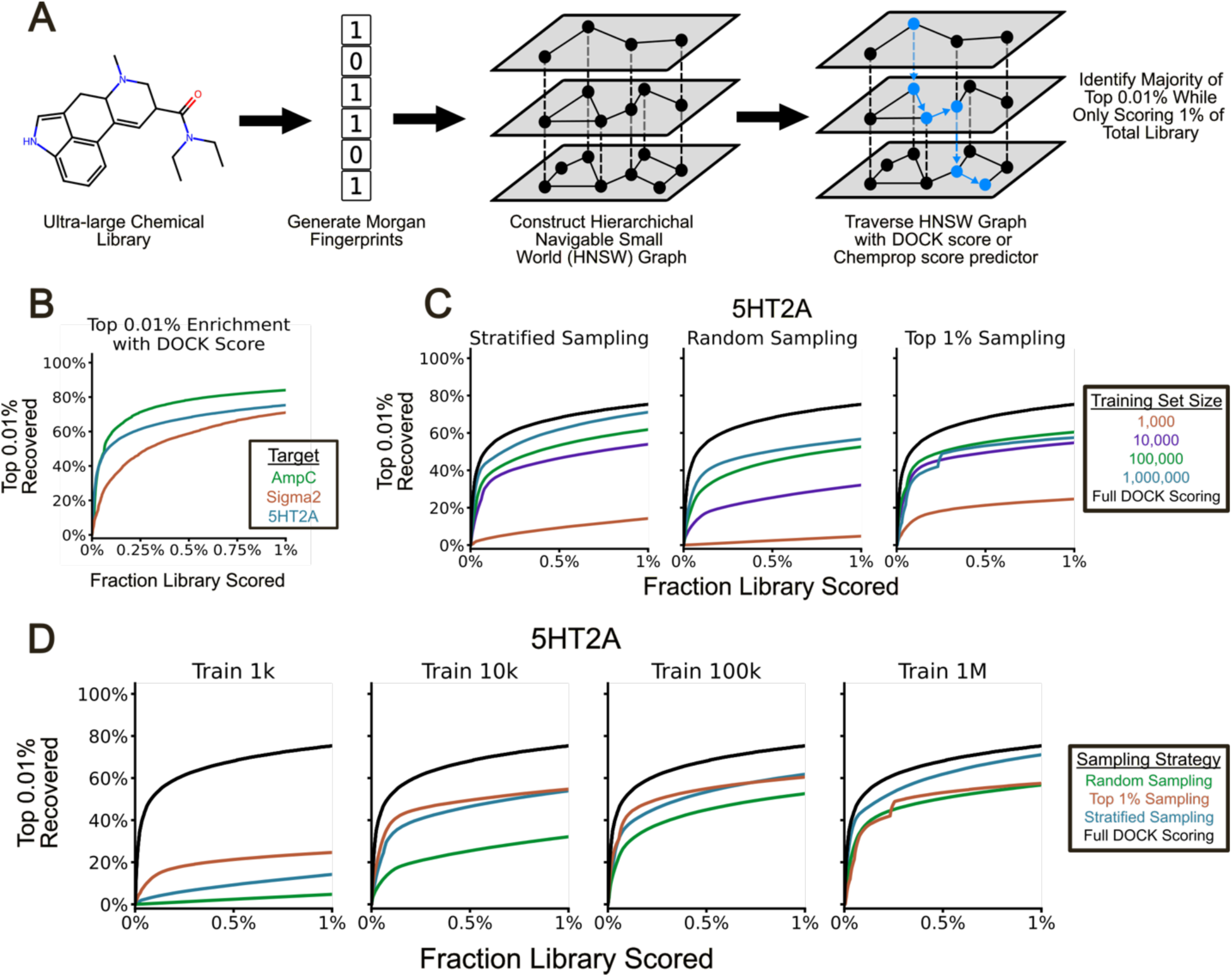
**(A)** Overview of Retrieval Augmented Docking (RAD). **(B)** Recall curves of the top 0.01% scoring molecules when evaluating a RAD-prioritized 1% of each library with the DOCK scoring function. **(C)** 5HT2A recall curves of the top 0.01% scoring molecules when evaluating a RAD-prioritized 1% of the library with the Chemprop models trained with different sampling strategies. The curves are averaged over five replicates. **(D)** 5HT2A recall curves of the top 0.01% scoring molecules when evaluating a RAD-prioritized 1% of the library with the Chemprop models trained with different training set sizes. The curves are averaged over five replicates.

On these systems, RAD found a large fraction of the top 0.01% scoring molecules while explicitly scoring only 1% of these billion molecule libraries (RAD deprioritizes the rest of the library as it traverses the graph). For the ∼1.3 billion molecule libraries, RAD found 84% of AmpC and 75% of 5HT2A top 0.01% scoring molecules (**Fig. 4B**). For the smaller 500 million molecule Sigma2 library, RAD found 71% of the top 0.01% scoring molecules, again while scoring a RAD-prioritized 1% of the total library (**Fig. 4B**).

We next asked whether the machine learning models trained to predict DOCK3.7/3.8 scores could effectively replace the DOCK scoring function in the context of RAD. With the best performing ML models, RAD using an ML scoring function found a comparable number of the top 0.01% scoring molecules as the full DOCK3.7/3.8 scoring function, but at much reduced computational cost due to the faster inference speed of machine learning models (∼1 sec/core to orient and score a molecule with DOCK3.8, versus ∼5 ms/core to do Chemprop ligand preparation and scoring). For example, RAD prioritization of 1% of the ∼1.3 billion 5HT2A library found 75% of the top 0.01% scoring molecules using the DOCK3.8 scoring function and 71% using the ML model trained on 1 million molecules with a stratified sampling strategy (**Fig. 4C**). We do note that the variance in RAD performance across replicates was larger than in the score prediction task (**Fig. S3**).

Here too, the models trained on larger datasets found more of the top 0.01% scoring molecules from a RAD-prioritized 1% of the library than those trained on smaller datasets. For instance, using the stratified sampling method with 5HT2A, RAD found 14%, 54%, 62%, and 71% of the top 0.01% scoring molecules among its top-prioritized 1% of the library with the models trained on 1,000, 10,000, 100,000, and 1,000,000 molecules, respectively (**Fig. 4C**). Models trained on larger datasets with the stratified sampling strategy continued to outperform those trained with random sampling or sampling only from the top 1% of scores (**Fig. 4D**). For example, on the 5HT2A library with models trained on 100,000 molecules, RAD found 62%, 60%, and 53% of the top 0.01% scoring molecules when screening a RAD-prioritized 1% of the library when using stratified sampling, top 1% sampling, and random sampling, respectively (**Fig. 4D**). These trends were also observed for the AmpC and Sigma2 datasets **(Fig. S4 and Fig. S5).**

## Discussion & Conclusions

There is much interest in training ML methods to reproduce docking results with a view to screening larger libraries faster, and naturally there is longstanding interest in improving scoring for docking and for virtual screening. Large benchmarks of docking results, both computational and experimental, would support these efforts but currently are unavailable to the community. Here we begin to develop an open access benchmark of docking and experimental results (lsd.docking.org). Three results merit emphasis. **First,** the scale of the benchmark may be appropriate to begin to support these goals. The docking screens are against eleven diverse targets, representing three unrelated enzymes, one transporter, and seven GPCRs (in three families), and over 6.3 billion molecules are scored. Docking scores are provided for all molecules successfully fit to the eleven targets, and for 10 million so too are docking poses. The results are presented in a format that we hope lends itself to machine reading. **Second**, the experimental activities are provided for 3,729 molecules across the eleven targets. These include both molecules that were experimentally active, as predicted by docking, and molecules that were docking false positives. For three targets, AmpC β-lactamase, Sigma2, and the dopamine D4 receptor, it includes the results not only of testing high-ranking molecules, but molecules across the scoring range—high-scoring, mediocre scoring, and poor scoring, all tested by experiment. As docking false positives can be informative both for scoring function optimization and ML training, these sets may be particularly useful. **Third,** in proof-of-concept studies, we investigated the sorts of ML model training and methods that the large-scale docking benchmarks might support. In particular, we trained ML models to predict a molecule’s DOCK score from its SMILES string and used these models as the scoring function for the HNSW method, itself drawn from large-scale vector database applications. In both cases, we found that increasing the training set size improved model performance. Despite some models obtaining high Pearson correlations between true scores and predictions, relatively few of the top 0.01% scoring molecules and true binders were in the top ML predictions, highlighting the pitfall of Pearson correlation as the sole evaluation metric. When applied in Retrieval Augmented Docking, the best-performing models were comparable to the full DOCK scoring function while much reducing computational cost.

Several caveats warrant mentioning. The scores reported here are all from DOCK3.7/3.8, a physics-based scoring function. This function has the advantage of allowing campaigns in the multi-billion molecule range, but the scores returned, though apparently correlated with hit-rate^1, 4, 39^, are offset from free energies of binding. Further, it represents only one of several widely used scoring functions. In the proof-of-concept ML training and optimization, a key limitation is the overlap between test sets and training sets. Although no molecule appeared in both the training and the test sets, we made no other effort to ensure a diverse or dissimilar test set, and model performance would likely decline when evaluated on a more diverse, out-of-distribution test set^50–52^. Since our goal was to showcase how the benchmarks *can* be used for optimization, rather than to develop specific optimized models, we hope the reader will see the results in this light. A similar limitation applies to the HNSW graphs, as molecules in the training sets may also appear in the graphs and be evaluated during traversal. However, even in the smaller ∼500 million Sigma2 library and with models trained on one million molecules, only 0.2% of the HNSW graph overlaps with the training set. While we do not expect this small overlap to significantly impact the results presented here, it remains a limitation of this study.

These cautions should not obscure the main observations from this study. The scale of docking and experimental results from recent virtual screening campaigns may support benchmarking sets for new ML models and methods. We provide the website lsd.docking.org to easily access these data and demonstrate their potential use in ML applications. As we continue to screen new targets at scale, we will upload new results to the website and welcome contributions from the community. It may particularly benefit from other larger library campaigns using other docking methods, which should be readily added to this resource.

## Methods

### Construction of the large-scale docking dataset (lsd.docking.org)

We identified directories on our computing cluster containing results from previously published large-scale docking campaigns run in our lab, with help from those who performed them. These directories included docking scores for all molecules in a campaign linked to ZINC IDs, which we used to query the ZINC database for the corresponding SMILES. We excluded a small fraction of ZINC IDs that no longer returned results due to updates of the ZINC database since the original campaigns. We combined the ZINC IDs, docking scores, and retrieved SMILES of all molecules in a campaign to produce the docking datasets available on lsd.docking.org. The directories also contained docking poses for molecules scoring better than a predefined threshold. We collected the docking poses for the top 500,000 scoring molecules without additional filtering and made them available on lsd.docking.org. For some campaigns, fewer than 500,000 poses were provided due to disk failures since the original campaign or due to fewer than 500,000 poses being saved during the original campaign. To complement these datasets, we uploaded the docking grids used to generate the poses and scores. Where available, we collected experimental results from the supplementary information of the corresponding publications. Otherwise, we located the experimental results with help from the lab member who performed the campaign and uploaded the results to lsd.docking.org (**Table S1**).

### Training Chemprop to predict DOCK scores

We evaluated the effects of three training set acquisition strategies and four training set sizes on the performance of ML models predicting DOCK scores from SMILES strings. The training set acquisition strategies included: random selection from the entire dataset, random selection from only the best 1% of molecules by DOCK score, and selecting 80% of the training data from the best 1% of molecules and the remaining 20% from the remainder of the dataset. We used training set sizes of 1,000, 10,000, 100,000, and 1,000,000.

We assessed model performance on test sets of 100,000,000 molecules using four metrics: the Pearson correlation between the predicted and actual DOCK scores across all test set molecules, the correlation only for the best 0.01% of test set molecules, the enrichment of the top 0.01% scoring molecules, measured by the log area under the curve (logAUC), which reflects recall as a function of ML ranking, and the enrichment of the actual new ligands discovered by experimental testing of top-ranked docking molecules. We applied these metrics to the docking results from three receptor campaigns: AmpC β-lactamase, 5HT2A, and Sigma2.

We randomly subsampled the AmpC, 5HT2A, and Sigma2 docking results from lsd.docking.org to create training sets. For each sampling strategy and training set size, we generated five independent training sets. To account for the validation set, we sampled an additional 20% of molecules. For instance, for a training set size of 10,000, we sampled an additional 2 000 molecules for the validation set, and for 100,000, we sampled an extra 20,000.

To minimize data leakage we split molecules based on their Bemis-Murcko scaffolds to assign them to the training and validation sets, ensuring no scaffold appeared in both sets. This approach produced training sets of approximately 1,000, 10,000, 100,000, and 1,000,000 molecules, with validation sets about 20% as large and containing no overlapping scaffolds.

We trained ML models with the Chemprop^40^ framework to predict DOCK scores from molecular SMILES strings. Chemprop, a PyTorch based framework, predicts molecular properties using message-passing neural networks on molecular graphs. We used the default parameters, featurizers, and settings. Specifically, we generated featurized molecular graphs from SMILES strings using the “SimpleMoleculeMolGraphFeaturizer,” messages were passed along directed bonds using “BondMessagePassing,” and the graph-level representation was obtained by averaging node-level representations with “MeanAggregation.” The learned representations had a dimensionality of 300, and a single-layer fully connected feed forward network predicted scores from the graph-level representation.

We trained the models to minimize the Root Mean Squared Error (RMSE) between predicted and true DOCK scores over 100 epochs. Training used the Adam optimizer with the Noam learning rate scheduler (initial, maximum, and final learning rates of 10^-4, 10^-3, and 10^-4, respectively). We used batch sizes of 64, batch normalization, and early stopping based on validation RMSE loss, with a patience of 5 epochs.

We gathered a test set for each receptor by randomly sampling 100,000,000 SMILES and DOCK scores for AmpC, 5HT2A, and Sigma2. Using the models trained with different sampling strategies and dataset sizes, we predicted DOCK scores for each molecule in the test set. After removing test set molecules included in a model’s training set, we calculated several metrics.

We calculated the Pearson correlation between predicted and actual DOCK scores for the entire test set and the top 0.01% scoring molecules in the test set. Next, we rank-ordered the ML predictions and plotted the fraction of the top 0.01% scoring molecules recovered against the ML ranking, using a log x-axis. We calculated the area under this curve (logAUC) to evaluate each model’s ability to enrich for the top 0.01% scoring molecules in its top predictions. This metric emphasizes practical utility in prospective screening, where only the top predictions are considered for visual inspection and testing.

### Hierarchical Navigable Small World (HNSW) graph construction and traversal

We demonstrated the utility of these billion-scale DOCK datasets by using them as benchmark sets for the chemical space exploration technique Retrieval Augmented Docking (RAD)^41^. RAD rapidly organizes chemical libraries into Hierarchical Navigable Small World (HNSW) graphs, which are multi-layered similarity-based structures built on the principles of small-world networks. The DOCK scoring function guides a best-first traversal of the HNSW graph, iteratively docking the neighbors of the best scoring nodes. Previous work found that RAD finds most of the top 0.01% scoring molecules while explicitly docking only a fraction of the library.

We constructed three HNSW graphs, one each for each receptor - AmpC, 5HT2A, and Sigma2 – using libraries from lsd.docking.org. The graphs contained 1,300 572,452, 1,300,808,533, and 468,639,652 molecules, respectively, randomly sampled from the corresponding docking results. We began by generating Morgan fingerprints with a radius of 2 and a length of 512 for all molecules. Using these fingerprints, we built the HNSW graphs with an *M* of 8, an *efConstruction* of 400, and the Tanimoto coefficient as the similarity metric. We then greedily traversed the HNSW graphs, as described in previous work^41^, using both the DOCK score and the predictions from the Chemprop models. We evaluated performance by measuring the fraction of the top 0.01% scoring molecules found while screening a RAD-prioritized 1% of the library.

## Author information

### Corresponding Authors

**Brian K. Shoichet** - Department of Pharmaceutical Chemistry, University of California, San Francisco, 94158, United States.

**John J. Irwin** - Department of Pharmaceutical Chemistry, University of California, San Francisco, 94158, United States.

## Authors

**Brendan W. Hall** - Department of Pharmaceutical Chemistry, University of California, San Francisco, 94158, United States.

**Tia A. Tummino** - Department of Pharmaceutical Chemistry, University of California, San Francisco, 94158, United States.

**Khanh Tang** - Department of Pharmaceutical Chemistry, University of California, San Francisco, 94158, United States.

## Author contributions

B.K.S. conceived the project. T.A.T. collected the data and prepared it for the database. K.T. made the website that hosts the database, guided by J.J.I. B.W.H trained the machine learning models and did the Retrieval Augmented Docking. B.K.S. and J.J.I. supervised the project. T.A.T. and B.W.H wrote the paper with input from all other authors, and with primary editing from B.K.S.

## Acknowledgements

We thank E. Fink, S. Gahbauer, J. Lyu, M. Rachman, I. Singh, R. Stein, and F. Liu for assistance with assembling the data that went into lsd.docking.org. We thank M. Castanon and O. Mailhot for help with algorithms for collecting ZINC22 SMILES for hundreds of millions of molecules.

## Notes

The authors declare the following competing financial interests: B.K.S. is co-founder of BlueDolphin LLC, Epiodyne, and Deep Apple Therapeutics, Inc., serves on the SRB of Genentech, the SABs of Schrödinger LLC and of Vilya Therapeutics. J.J.I. is a co-founder of BlueDolphin LLC and of Deep Apple Therapeutics. The authors declare no other competing interests.

## Data and Software Availability

All docking results, docking poses, and experimental data are made freely available at lsd.docking.org. Open-source code for performing Retrieval Augmented Docking is available at https://github.com/keiserlab/rad. Open-source code for replicating the Chemprop model training, inference, and integration with RAD is available at https://github.com/bwhall61/lsd.

## Abbreviations Used

GPCR: G Protein-Coupled Receptor
ML: Machine Learning
RAD: Retrieval Augmented Docking
HNSW: Hierarchical Navigable Small World
logAUC: Logarithmic Area Under the Curve
LSD: Large-Scale Docking
SMILES: Simplified Molecular Input Line Entry System
5HT2A: 5-hydroxytryptamine (serotonin) receptor 2A
AmpC: AmpC β-lactamase
RMSE: Root Mean Squared Error

## Table of Contents Graphic

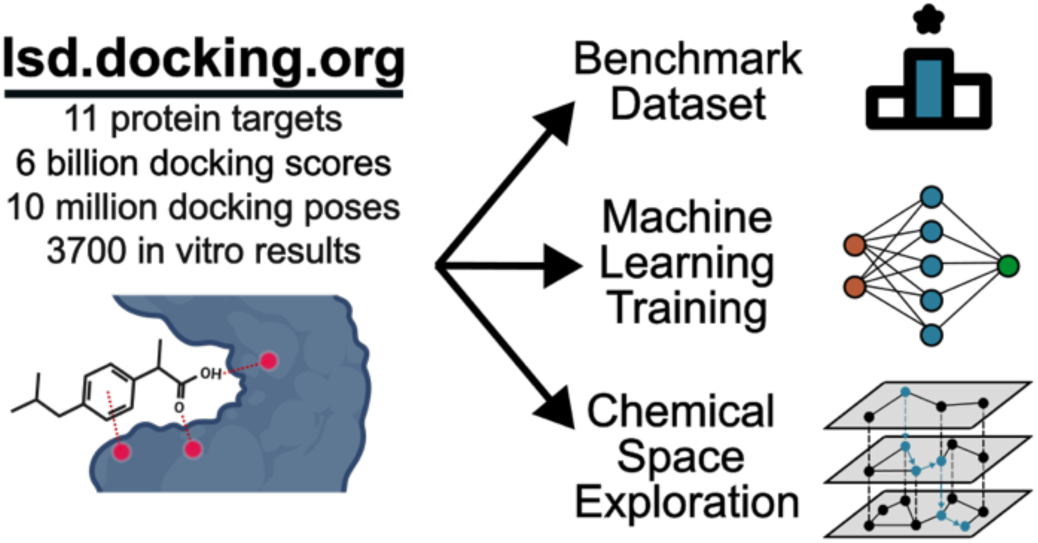

