## Supporting Information for "A database for large-scale docking and experimental results"

### ASSOCIATED CONTENT

& Brian K. Shoichet[1,*]

[1]Department of Pharmaceutical Chemistry, University of California, San Francisco, San Francisco, CA 94158, USA

† These authors contributed equally.

| Target | Docking Scores | Docking Poses | Experimental Results |
|---|---|---|---|
| **Alpha2AR** | | | |
| Fink_2022 | | | |
| Fragments1 | 7,907,562 | 161,330 | 35 |
| Fragments2 | 9,168,485 | 500,000 | 42 |
| Leadlike | 13,442,764 | 500,000 | 5 |
| **AmpC** | | | |
| Lyu_2019 | 99,459,561 | 500,000 | 44 |
| Liu_2025 | 1,468,863,655 | 500,000 | 1521 |
| **CB1R** | | | |
| Tummino_2024 | 18,992,691 | 500,000 | 46 |
| **D4** | | | |
| Lyu_2019 | 138,312,677 | 500,000 | 552 |
| **EP4R** | | | |
| Gahbauer_2023 | | | |
| Screen1 | 344,092,434 | 500,000 | 40 |
| Screen2 | 36,974,635 | 500,000 | 31 |
| **MPro** | | | |
| Fink_2023 | | | |
| *Covalent* | | | |
| Aldehydes | 1,392,177 | 500,000 | 27 |
| Ketoamides | 393,075 | 0 | 15 |
| Nitriles1 | 537,016 | 0 | 6 |
| Nitriles2 | 535,968 | 0 | 12 |
| *Noncovalent* | | | |
| Screen1_ZINC15 | | | |
| bigger | 107,486,710 | 500,000 | 89 |
| leadlike | 219,305,079 | 500,000 | 105 |
| Screen2_ZINC22 | 778,517,250 | 499,990 | 139 |
| **MT1R** | | | |
| Stein_2020 | 40,376,489 | 499,999 | 38 |
| **NSP3_Mac1** | | | |
| Gahbauer_2022 | | | |
| Everted_ZINC22 | 57,437,478 | 500,000 | 56 |
| FRESH_anions_Feb2020 | 15,957,174 | 500,000 | 9 |
| MOD_Leadlike1 | 316,505,043 | 500,000 | 78 |
| MOD_Leadlike2 | 239,255,129 | 500,000 | 23 |
| REAL_anions | 37,556,136 | 500,000 | 14 |
| Schuller_2021 | | | |
| InMan | 17,362 | 10,693 | 10 |
| InStock_Fragments | 696,092 | 345,695 | 8 |
| MOD_Fragments | 19,130,798 | 500,000 | 42 |
| **SERT** | | | |
| Singh_2023 | 246,614,514 | 404,706 | 13 |
| **Sigma2** | | | |
| Alon_2021 | 468,639,651 | 500,000 | 506 |
| **5HT2A** | | | |
| Lyu_2024 | 1,630,264,067 | 500,000 | 223 |

**Table S1.** Overview of the docking screens included on lsd.docking.org and the number of docking scores and experimental results provided for each screen.

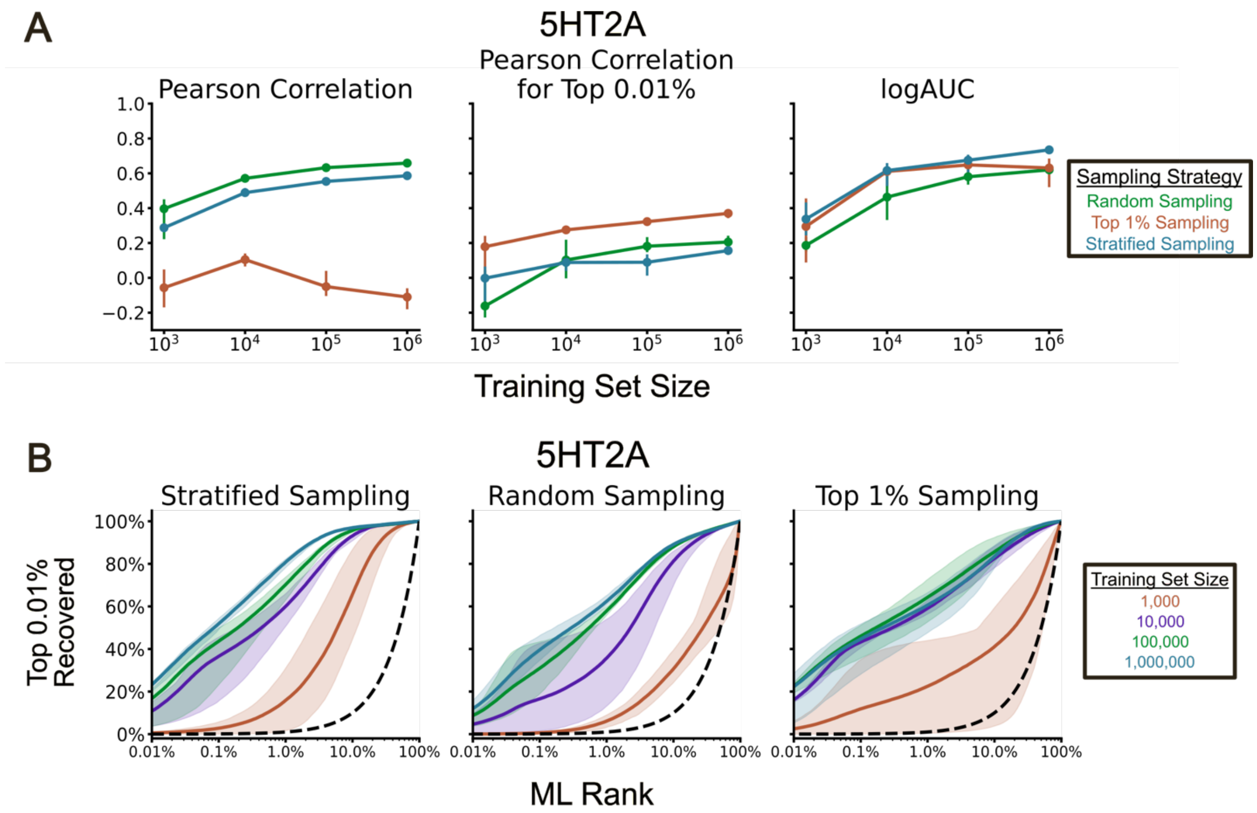


**Figure S1. (A)** Effects of increasing the training set size on the three evaluation metrics for the 5HT2A receptor. The bars represent the minimum and maximum performance across five replicates. **(B)** 5HT2A recall curves of the top 0.01% scoring molecules for the three sampling strategies using increasing training set sizes. The shaded region represents the minimum and maximum performance across five replicates. The dashed black line represents a random predictor.

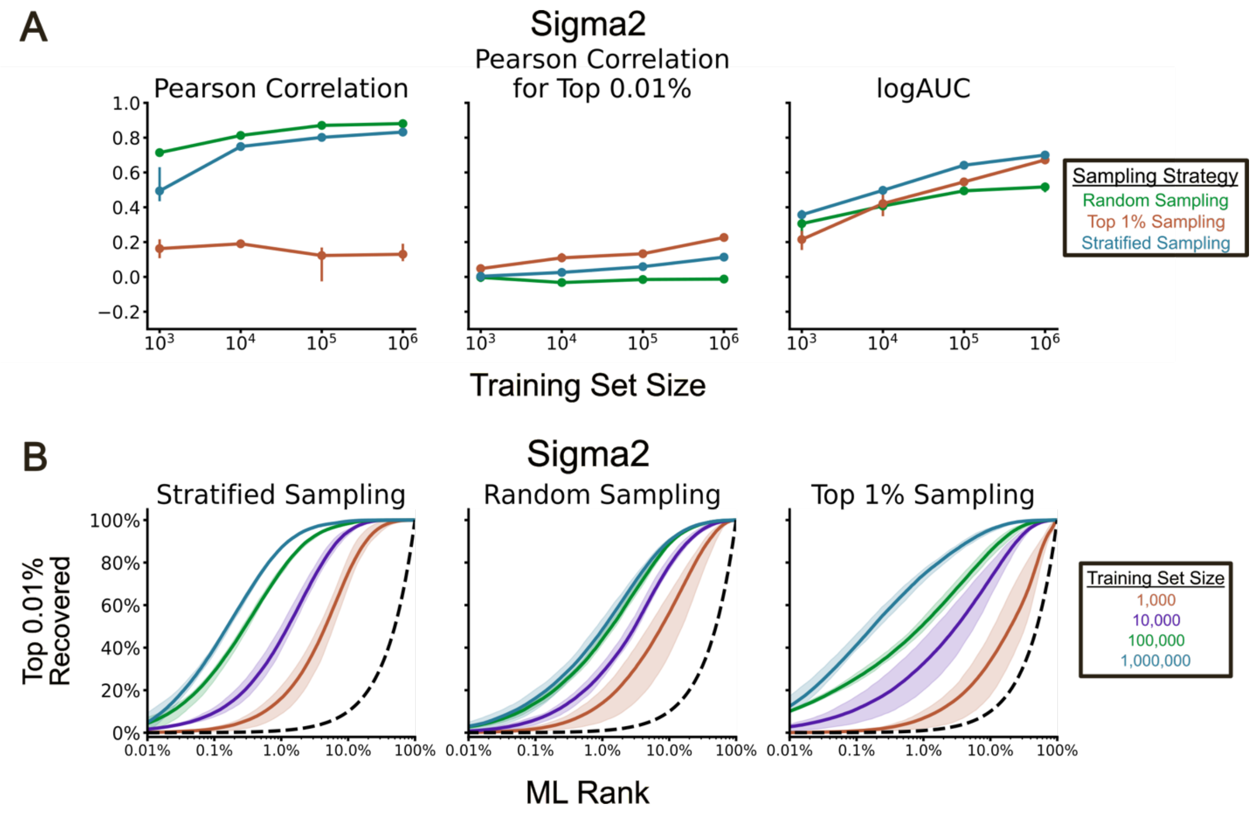


**Figure S2. (A)** Effects of increasing the training set size on the three evaluation metrics for the Sigma2 receptor. The bars represent the minimum and maximum performance across five replicates. **(B)** Sigma2 recall curves of the top 0.01% scoring molecules for the three sampling strategies using increasing training set sizes. The shaded region represents the minimum and maximum performance across five replicates. The dashed black line represents a random predictor.

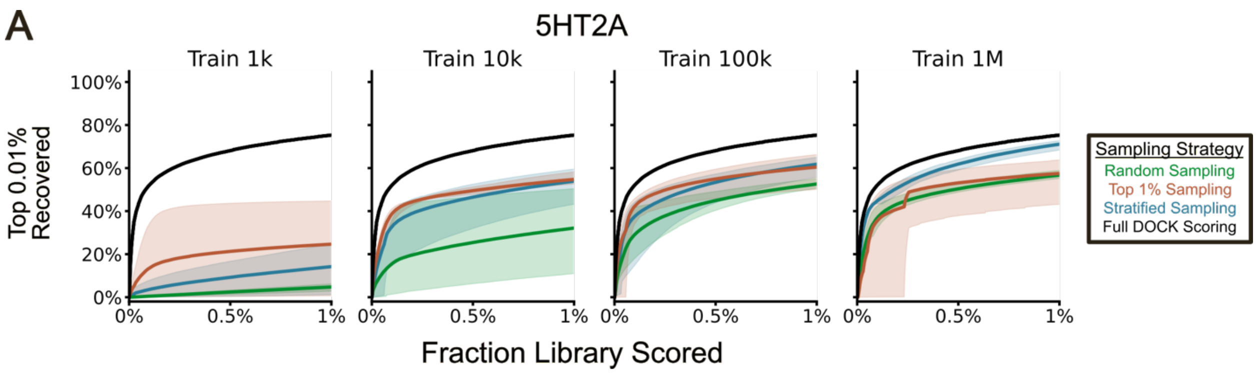


**Figure S3.** 5HT2A recall curves of the top 0.01% scoring molecules when evaluating a RAD-prioritized 1% of the library with the Chemprop models trained with different training set sizes. The shaded regions represent the minimum and maximum performance over five replicates.

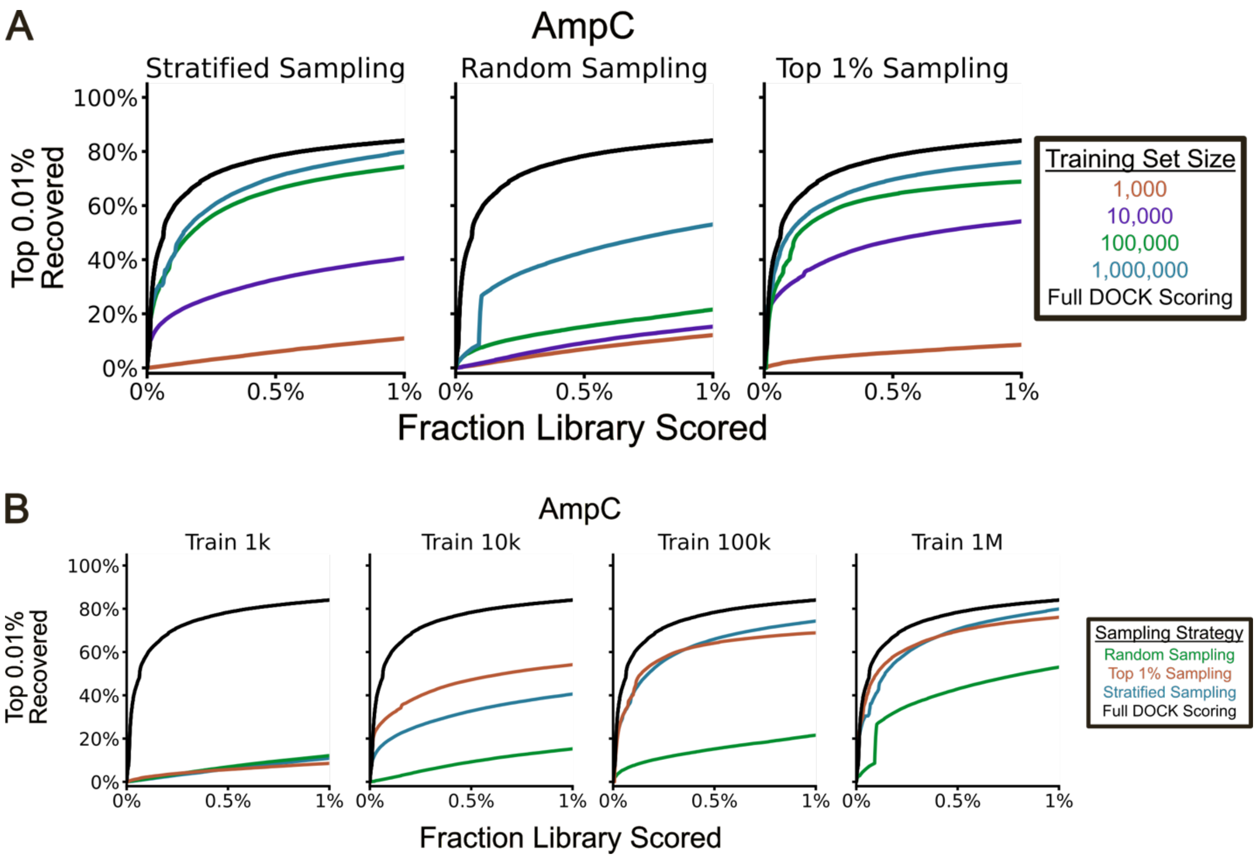


**Figure S4. (A)** AmpC recall curves of the top 0.01% scoring molecules when evaluating a RAD-prioritized 1% of the library with the Chemprop models trained with different sampling strategies. The curves are a single replicate. **(B)** AmpC recall curves of the top 0.01% scoring molecules when evaluating a RAD-prioritized 1% of the library with the Chemprop models trained with different training set sizes. The curves are a single replicate.

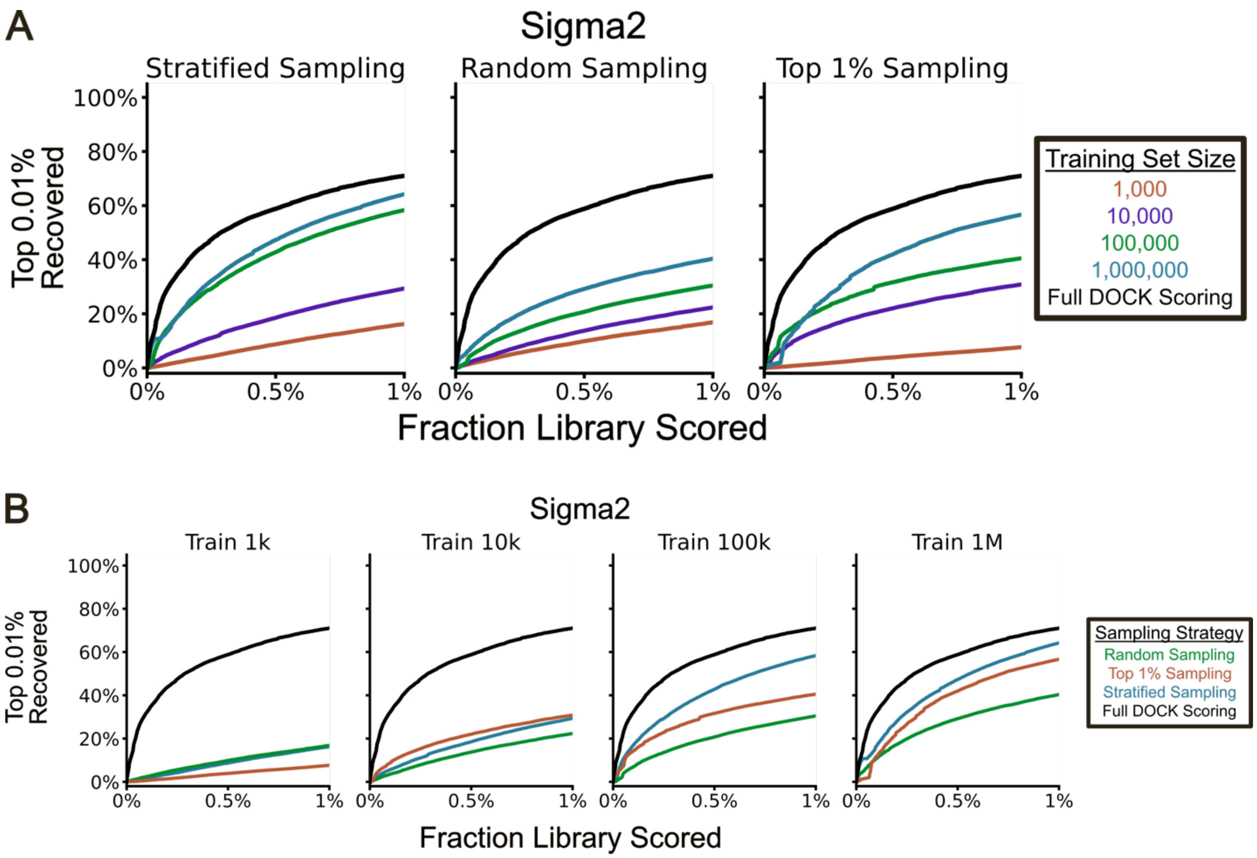


**Figure S5. (A)** Sigma2 recall curves of the top 0.01% scoring molecules when evaluating a RAD-prioritized 1% of the library with the Chemprop models trained with different sampling strategies. The curves are a single replicate. **(B)** Sigma2 recall curves of the top 0.01% scoring molecules when evaluating a RAD-prioritized 1% of the library with the Chemprop models trained with different training set sizes. The curves are a single replicate.
